# High content screening and computational prediction reveal viral genes that suppress innate immune response

**DOI:** 10.1101/2021.12.14.472572

**Authors:** Tai L. Ng, Erika J. Olson, Tae Yeon Yoo, H. Sloane Weiss, Yukiye Koide, Peter D. Koch, Nathan J. Rollins, Pia Mach, Tobias Meisinger, Trenton Bricken, Timothy Z. Chang, Colin Molloy, Jérôme Zürcher, Timothy J. Mitchison, John I. Glass, Debora S. Marks, Jeffrey C. Way, Pamela A. Silver

## Abstract

Suppression of the host innate immune response is a critical aspect of viral replication. Upon infection, viruses may introduce one or more proteins that inhibit key immune pathways, such as the type I interferon pathway. However, the ability to predict and evaluate viral protein bioactivity on targeted pathways remains challenging and is typically done on a single virus/gene basis. Here, we present a medium-throughput high-content cell-based assay to reveal the immunosuppressive effects of viral proteins. To test the predictive power of our approach, we developed a library of 800 genes encoding known, predicted, and uncharacterized human viral genes. We find that previously known immune suppressors from numerous viral families such as *Picornaviridae* and *Flaviviridae* recorded positive responses. These include a number of viral proteases for which we further confirmed that innate immune suppression depends on protease activity. A class of predicted inhibitors encoded by *Rhabdoviridae* viruses was demonstrated to block nuclear transport, and several previously uncharacterized proteins from uncultivated viruses were shown to inhibit nuclear transport of the transcription factors NF-κB and IRF3. We propose that this pathway-based assay, together with early sequencing, gene synthesis, and viral infection studies, could partly serve as the basis for rapid *in vitro* characterization of novel viral proteins.

**IMPORTANCE:** Infectious diseases caused by viral pathogens exacerbate healthcare and economic burdens. Numerous viral biomolecules suppress the human innate immune system, enabling viruses to evade an immune response from the host. Despite our current understanding of viral replications and immune evasion, new viral proteins, including those encoded by uncultivated viruses or emerging viruses, are being unearthed at a rapid pace from large scale sequencing and surveillance projects. The use of medium- and high-throughput functional assays to characterize immunosuppressive functions of viral proteins can advance our understanding of viral replication and possibly treatment of infections. In this study we assembled a large viral gene library from diverse viral families and developed a high content assay to test for inhibition of innate immunity pathways. Our work expands the tools that can rapidly link sequence and protein function, representing a practical step towards early-stage evaluation of emerging and understudied viruses.

## INTRODUCTION

Pathogenic viruses (*e.g*. Ebola, HIV, SARS-CoV-2) continue to pose public health threats and cause economic disruptions worldwide. The human innate immune system has evolved multiple signaling pathways, including the type I interferon pathway to defend against viral infections. These pathways use nucleic acid receptors to trigger timely immune responses, including the expression of proteins that halt viral replication and production of interferon that activates the JAK/STAT signaling pathway (1, 2). Viruses have evolved several ways to evade these mechanisms, such as binding or degrading proteins in these pathways, molecular mimicry, or modulating host gene expression (3–6). Identifying viral proteins that block immune signaling leads to potential drug targets and a significantly improved understanding of viral replication.

Despite substantial progress in our knowledge of viral pathogenicity and immune evasion, a standardized and consistent study for rapidly investigating many different viruses remains underexplored due to challenges such as optimizing virus cultivation and defining cell type. Furthermore, next generation sequencing and proteomics have provided an abundance of uncharacterized viral proteins, with many generically annotated as ‘nonstructural proteins’ or ‘hypothetical proteins’ (7). Even for functionally validated proteins, annotations inferred across species may prove to be inaccurate without experimental validations. Some viral enzymes, such as viral proteases, may have unidentified moonlighting roles as immune suppressors (8). These uncharacterized sequences are expected to continue to grow massively from large-scale virus collection and surveillance projects (9). Altogether, these challenges limit our understanding of viral pathogenicity, and consequently, treatment of viral infections.

We envision a multiprong approach using a suite of assays that can rapidly identify different functions of unknown viral proteins. Such screens could complement viral infection studies pursued by academic labs or dedicated government facilities for viral surveillance. Ultimately these assays will further our knowledge of immune evasion by viruses. We explored this concept by starting with the development of a medium-throughput, microscopy-based immune assay in fibroblast BJ-5ta cells. We assembled a library of 605 viral genes encoded by viruses from 31 viral families, including 536 sequences with unannotated immunosuppressive function. We tested an additional 195 coronavirus genes during the COVID-19 pandemic. Our assay identified many inhibitors, some of which were previously reported, some with immunosuppressive function that was inferred from sequence similarity or Pfam homology, and some proteins with unsuspected potential for immune suppression.

## RESULTS

### Viral gene library construction

First, we aimed to develop a library of viral genes that would further our understanding of human viruses (**Figure 1**). We envisioned that our library should contain known immune inhibitors, homologues of immune inhibitors, and uncharacterized proteins. We also focused on human and insect host viruses to highlight relevance to diseases, as well as understudied viruses from diverse viral families. 6,000 protein sequences were collected from GenBank reference genomes of the 1,688 human viruses in VirusHostDB (10). To sample diverse proteins spanning this set, we clustered at >20% sequence identity and >80% coverage using CD–HIT (11, 12), resulting in 1,975 clusters. Well-conserved viral proteins such as capsids and replication enzymes were collected in large clusters of up to over 100 members, whereas hundreds of sequences had no close relatives in other reference genomes. To increase the chances of identifying immune suppressors, we focused on smaller genomes (< 35 genes). We also excluded likely integral membrane proteins, as those may not properly fold in the absence of other viral proteins. We also excluded large polypeptides that are typically cleaved in the context of a viral infection.

**Figure 1:**
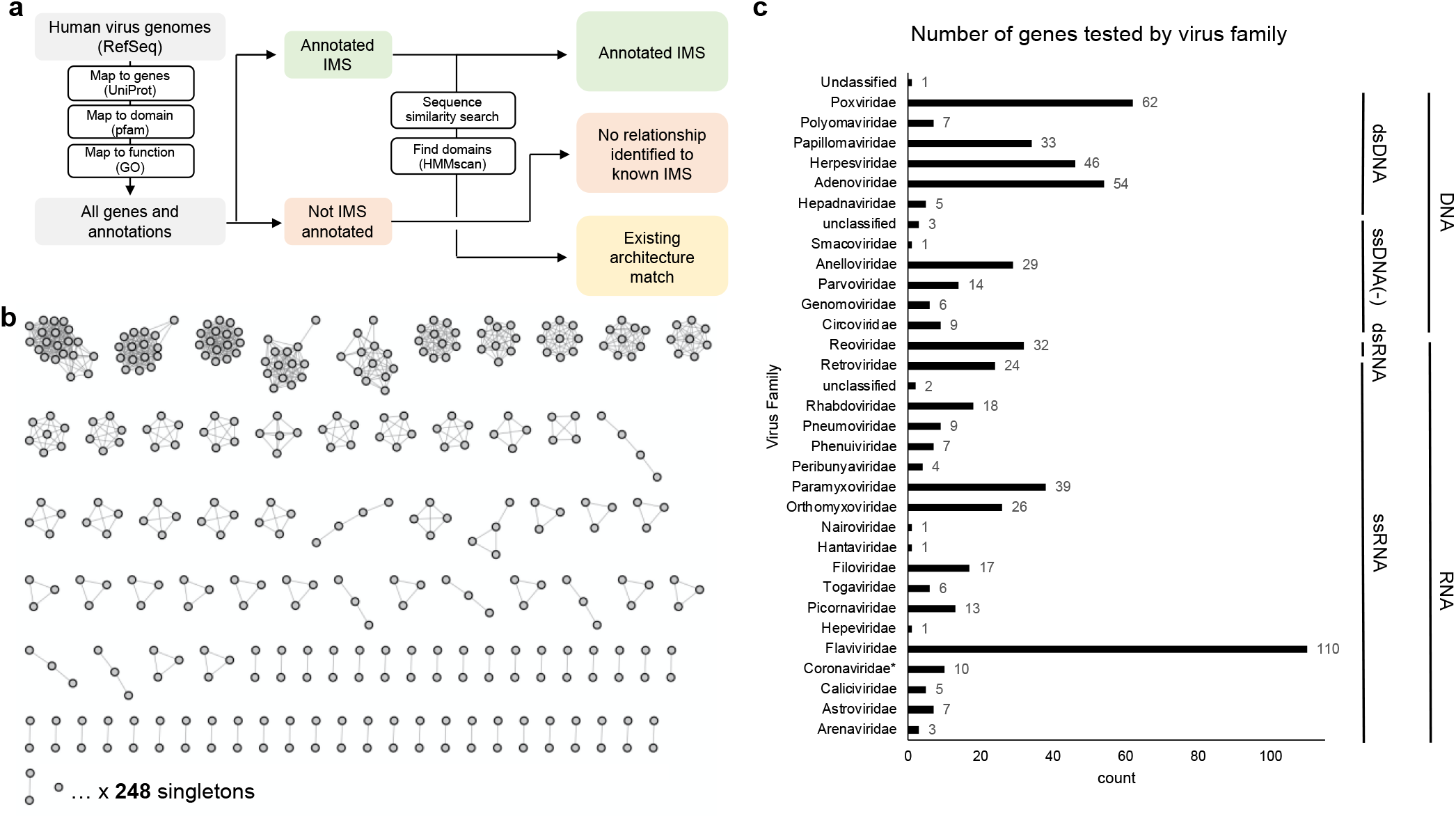
Assembly of a viral gene library to test in immune suppression screens. **(a)** Overview of the bioinformatic workflow to generate a list of viral proteins for testing. We designated viral genes as 1) immunosuppressive (IMS) by gene ontology (GO) or Pfam search, 2) predicted IMS based on sequence similarity or permissive hmmscan, or 3) uncharacterized viral proteins. **(b)** Sequence similarity network of 605 viral proteins to test clusters of sequence-related proteins and singletons. **(c)** Distribution of 605 genes by viral family. 195 additional genes from Coronaviridae were tested during the COVID-19 pandemic and are not included in this figure.

To collect viral genes known and predicted to inhibit the innate immune system, we searched for gene ontology (GO) annotations (13, 14) down the tree of high-level terms for virus suppression of host innate immune responses and apoptosis (GO:0039503, GO:0052170, GO:0052309, GO:0019050). Proteins annotated with these terms, derived from human and insect-infecting viruses, and confirmed protein expression reported in UniProt, serve as the known innate immune suppressors (**Data Set S1**). Sequences without these gene ontology annotations but that have 20% pairwise alignment identity to the known inhibitors form the group of viral proteins with predicted immunosuppressive activities. Functionally related genes with low sequence similarity were identified by deeper models that capture sequence variation across proteins with similar functions (*e.g*. sharing a family or domain); Pfam is a curated database of Hidden Markov Models (HMMs) capturing that information (15). We used these HMMs to categorize protein regions as functionally related to known families with high-confidence categorizations that are available on Pfam and UniProt. Lower confidence domain categorizations can be considered hypothetical and are potential candidates for functional characterization. Overall, we can hypothesize gene functions that are highly distant in sequence similarity from shared domains. Searching our viral protein sequence database (both annotated and unannotated immunosuppressive genes by GO) resulted in 42 proteins as positive hits and 243 predicted inhibitors identified with a more permissive hmmscan.

Overall, our library of 605 genes contains 69 viral immune inhibitors with GO annotations containing immune suppressive (IMS) terms, 158 viral genes with 20% sequence similarity to the IMS genes. Additionally, this library contains 42 known inhibitors determined by containing an immunosuppressive domain by Pfam classification and 243 predicted inhibitors using permissive hmmscan (**Data Set S1**). 358 genes in this library are not predicted to be immunosuppressive. In total, we tested 269 proteins from DNA viruses, 335 proteins from RNA viruses (including retroviruses), and 1 protein from an unclassified hudisavirus (**Figure 1C**). The COVID-19 pandemic occurred in the middle of our screening effort described below, which prompted us to test 195 viral genes encoded by the coronaviruses SARS-CoV-2, SARS-CoV, MERS-CoV, hCoV-229E, hCoV-NL63, hCoV-OC43, and hCoV-HKU1 (**Data Set S2**).

### High content screening assay development

To assay the gene library for suppression of human innate immunity, we developed a high-content, medium-throughput assay to reveal the effects of individual viral proteins on the type I interferon pathways. The three main signaling cascades tested are the TLR3 (toll-like receptor 3)-, cGAS–cGAMP–STING-, and JAK/STAT-mediated pathways. These three signaling axes respectively respond to foreign endosomal RNA (i, **Figure 2a**), cytosolic DNA (iii), and interferon (IFN, iv). When the cell senses foreign DNA and RNA via pattern recognition receptors, multiple signaling cascades lead to nuclear translocation of cytosolic pro-inflammatory transcription factors NF-κB, IRF3, and pSTAT, which will produce antiviral responses. We chose these signaling axes because the pathways and numerous viral inhibitors have been well-studied in the literature (1, 2). We developed conditions in which the nuclear/cytoplasmic localization of IRF3 and NF-κB could be simultaneously visualized using non-cross-reacting primary and secondary antibodies (16).

**Figure 2:**
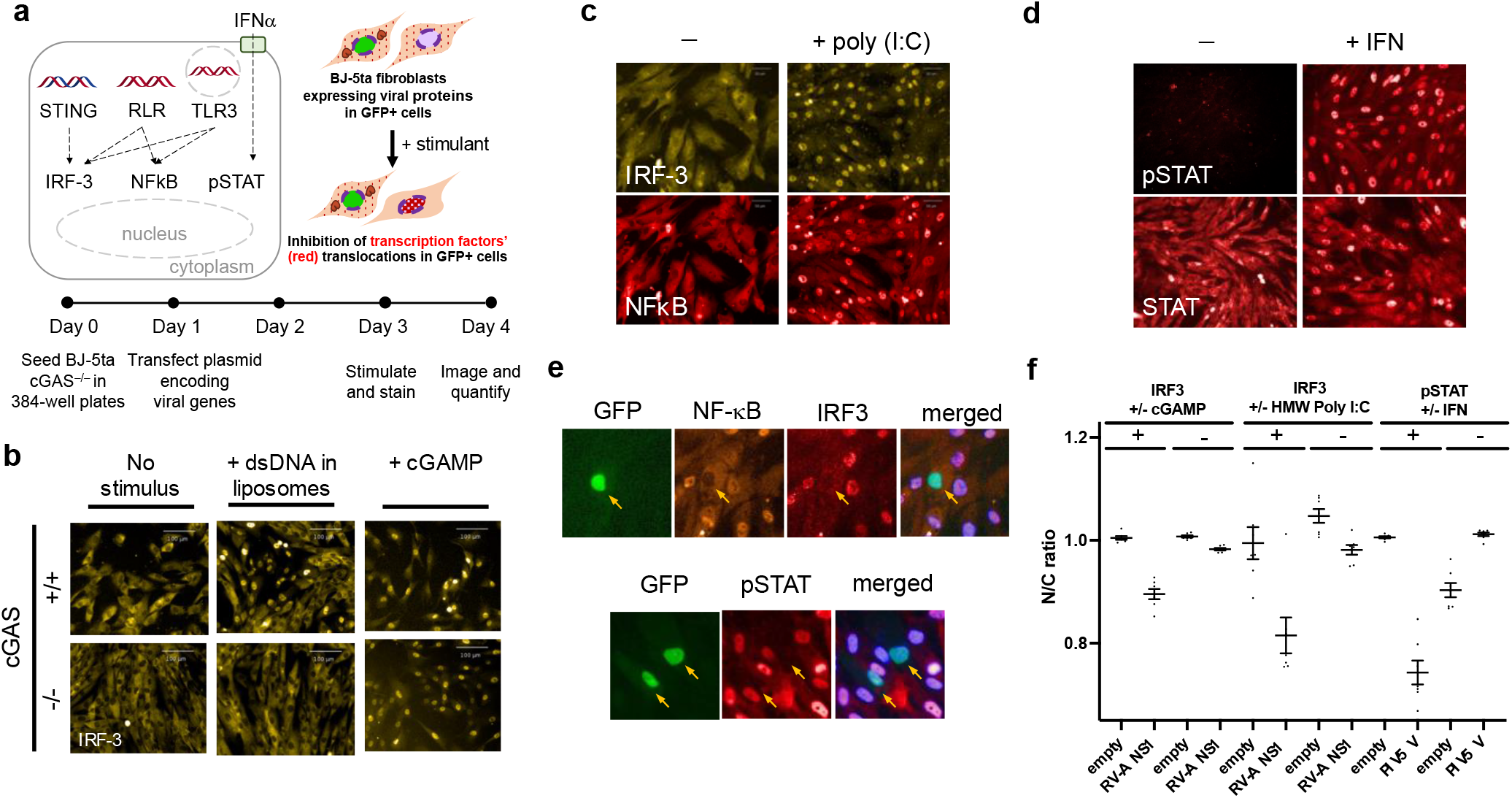
Activation and suppression of antiviral innate immune pathways assayed in fibroblasts via medium-throughput fluorescence microscopy. **(a)** Innate immune system signaling pathways that respond to viral infection. Top left: Transcription factors IRF3 and NF-κB are activated by the presence of nucleic acids in an inappropriate cellular compartment, signifying viral infection. Together, IRF3 and NF-κB activate expression of α and β interferons (IFNs), which are secreted and locally stimulate the JAK/STAT pathway. IRF3, NF-κB and pSTAT1 are all nuclear translocated proteins during signaling. Viruses often encode proteins that disrupt these pathways, by directly or indirectly inhibiting nuclear translocation, or by causing degradation of the transcription factor. Top right: The pathways can be initiated *in vitro* by addition of cGAMP (a second messenger), extracellular dsRNA, or by IFNα. Bottom: Experimental workflow for testing virus genes for modulation of the IRF3, NF-κB, and pSTAT1 signaling pathways. BJ-5ta (cGAS^−^) cells are transiently transfected with a viral gene expression vector, treated with various stimuli, and then fixed, stained with antibodies against IRF3, NF-κB, and/or pTyr701-STAT1, incubated with secondary antibodies, and then imaged. **(b)** Knockout of cGAS prevents transfection-mediated stimulation of IRF3 translocation while allowing downstream activation of IRF3 via cGAMP. Double-stranded DNA in the cytoplasm activates cGAS to create the cyclic dinucleotide cGAMP, which then acts on STING to activate IRF3. Parental cGAS+ BJ-5ta cells show nuclear IRF3 translocation in response to transfected DNA and exogenous cGAMP, while BJ-5ta cells with a CRISPR cGAS knockout show an IRF3 response only after treatment with cGAMP. This allowed us to assay elements of the STING/IRF3 pathway without interference by transfected DNA. **(c)** BJ-5ta cells treated with poly(I:C) show translocation of IRF3 and NF-κB into the nucleus. **(d)** IFNα treatment caused translocation of cytoplasmic nuclear STAT1 and pSTAT1 into the nucleus. For this study, we chose to only stain for pSTAT1. **(e)** Images of BJ-5ta cells that were co-transfected with plasmids encoding GFP and rotavirus A (RV-A) nonstructural protein (NS) 1 and stained for NF-κB and IRF3. Cells were also co-transfected with encoding GFP and parainfluenza 5 (PIV5) V protein and stained for pSTAT1. Only transfected cells (arrows) show inhibition of nuclear localization or formation of phospho-STAT1. **(f)** Quantitative results for nuclear-to-cytoplasmic (N/C) ratio for IRF3 and pSTAT1 for BJ-5ta cells expressing no viral protein, rotavirus A NS1, or PIV5 V protein. Some cells expressing these proteins were treated with cGAMP, high molecular weight poly(I:C), or IFNα to activate the immune signaling axes of interest. Viral inhibition of transcription factor translocation resulted in lower N/C ratio in the presence of stimuli.

All viral genes were synthesized and placed into a plasmid vector to promote constitutive expression upon transfection. We chose non-transformed, tert-immortalized BJ-5ta fibroblasts as our host cell type, as these cells are amenable to imaged-based screening and high-throughput screens (16). Furthermore, BJ-5ta fibroblast encode relatively intact innate immune pathways compared to other cell types (*e.g*. HEK293T cells express low levels of STING (17) and TLR3 (18)). Since DNA transfection alone stimulates the cGAS-STING pathway (**Figure 2e**), we generated a CRISPR knockout of cGAS (BJ-5ta (cGAS^−^), Methods). In this cell line, DNA transfection-mediated stimulation of IRF3 nuclear transport was essentially undetectable (**Figure 2b**).

To assess the effect of a particular gene on innate immune signaling pathways, BJ-5ta (cGAS^−^) cells (**Figure 2a**) were seeded in 384-well plates and co-transfected with a viral gene expression vector and a GFP expression plasmid. Extracellular poly(I:C) or cGAMP were used to stimulate signaling via TLR3 or STING pathways respectively. These stimuli caused localization of IRF3 and/or NF-κB from the cytoplasm to the nucleus (**Figures 2d** and **2e**). BJ-5ta (cGAS^−^) cells were also treated with IFNα to initiate STAT1 phosphorylation and nuclear translocation (**Figure 2f**). Two days after stimulation, cells were stained with either a mixture of NF-κB and IRF3 or with anti-pSTAT1 (phospho-pSTAT1 Tyr701) antibodies. Based on the timing of the induced response, cells were fixed, stained with antibodies directed against relevant transcription factors, and imaged. Fields of cells were scored by automated image processing for expression of a GFP transfection marker and the nuclear/cytoplasmic distribution of NF-κB, IRF3, or pSTAT1 (**Figure 2c, d, e**). ‘Hits’ were identified based on inhibition strengths of cells transfected with a viral protein-encoding vector compared to an empty vector control (see Methods and Supplemental Information). Using this assay, we tested rotaviral A (RV-A) nonstructural protein 1 (NS1), which induces IRF3 degradation (19). Significant inhibitory responses of IRF3 translocation were observed when cells were treated with cGAMP or poly(I:C) (**Figure 2f**). We also tested parainfluenza virus (PIV) 5 V protein, a known JAK/STAT pathway inhibitor, and observed strong inhibition of pSTAT translocation when the BJ-5ta cells were treated with IFNα (20). Therefore, this workflow is suitable for screening our viral gene library for immunosuppressive functions.

### High content imaging assays identified known and predicted viral inhibitors

Using the described assays, we tested 605 viral genes from our library (**Figure 1c**) and an additional 195 coronaviral genes we obtained during the Covid-19 pandemic. Applying a ‘stringent’ cutoff (p<0.05) by comparison to the no stimulus control (**Figures S1–3**) resulted in clear enrichment of 79 viral proteins that inhibited transcription factor translocation (**Figure 3a**, **Data Set S4**). Only a few proteins that strongly inhibited NF-κB translocation in the cGAMP-stimulated assay were obtained (**Figure 3a, Figure S2b**). Overall, we obtained more IRF3 translocation inhibitors than NF-κB translocation inhibitors in the STING- and TLR3-mediated pathways, and our assays revealed more pathway-specific inhibitors than proteins that inhibit multiple signaling axes (**Figure 3b**).

**Figure 3:**
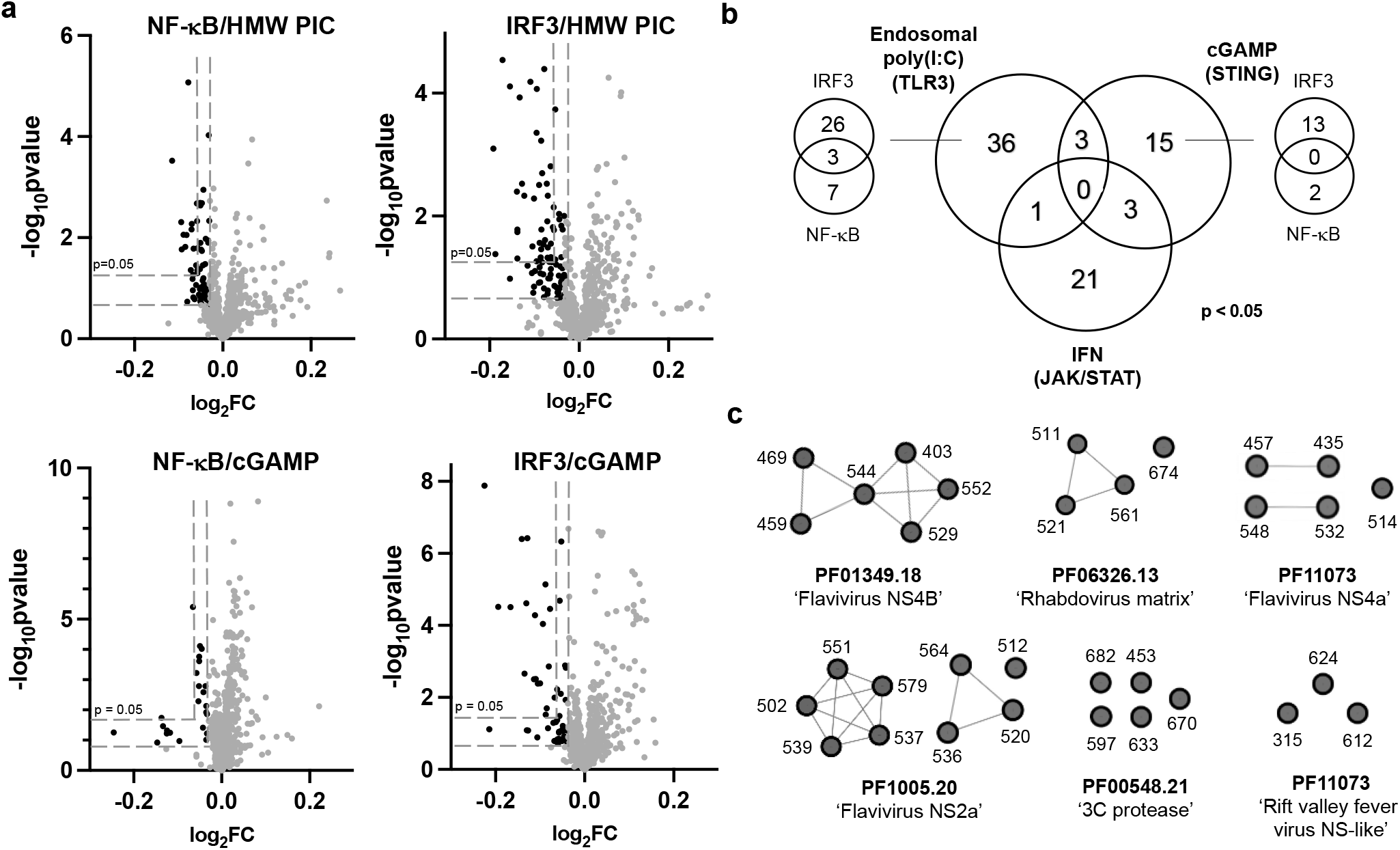
**(a)** Volcano plots highlight hits with stringent (p<0.05) and permissive cutoffs. 800 genes (including the 195 coronavirus genes) are plotted. Cutoff values for log_2_fold change (log_2_FC) and p-values were determined by comparing the data to the corresponding no treatment controls, which do not result in robust nuclear translocation of transcription factors. Results for no treatment controls and IFNα-treated cells are depicted in **Figures S1–3**. Raw source data is provided in **Data Set S3**. **(b)** Venn diagram depicting 79 stringent hits (p<0.05) among 800 viral genes across the four assays. For the endosomal (HMW poly(I:C))- and STING (cGAMP)-stimulated pathways, the additional Venn diagrams report the number of viral genes that inhibited IRF3 and/or NF-κB. **(c)** Examples of sequence-related positive innate immune inhibitors, grouped by Pfam domains, found in our screen and/or among inhibitors reported in the literature. The full list of permissive hits can be found in **Data Set S5** and the full sequence similarity network file of permissive hits can be found in Source Data Files.

The list of 79 viral proteins included numerous known inhibitors of innate immunity (**Data Set S4**). Specifically, 3 proteins were annotated with our selected immunosuppressive (IMS) gene ontology terms, and 16 proteins that were related to positive IMS genes by sequence similarity. 7 proteins were identified that encode protein domains known to inhibit immune function and 18 viral proteins inferred from hmmscans of known inhibitors. Two viral proteins, flaviviral NS4B and parvoviral VP1, were hits that contain domains with homology to known human proteins. Examples of previously reported inhibitors included cowpox viral poxin, which strongly inhibited IRF3 translocation when the cells are stimulated with cGAMP but not other stimuli. This observation is consistent with poxin nuclease’s cGAMP-cleaving activity (21). W protein from Nipah henipavirus inhibited nuclear translocation of IRF3 when stimulated with HMW poly(I:C). This protein has been demonstrated to inhibit phosphorylation of IRF3 (22). Other strong hits include several phenuiviral nonstructural proteins (NSs), picornarviral ‘3C’ proteases (3C^pro^), and rhabdoviral matrix proteins (M) (**Figure 3b**). NSs proteins from phenuivirus, specifically sandfly fever Sicilian virus and heartland virus, are known to block the DNA-binding domain of IRF3 (23, 24). Our assays demonstrated that NSs from sand fever Turkey virus and heartland virus both inhibited IRF3 translocation when stimulated with poly(I:C). Moreover, heartland virus NSs inhibited cGAMP-stimulated IRF3 translocation while the protein encoded by sand fever Turkey virus inhibited to a lesser extent. Finally, we identified 14 coronavirus genes as immune inhibitors, including SARS-CoV-2 Orf3a (IRF3, poly(I:C)-treated), Nsp10 (IRF3 and pSTAT, poly(I:C)- and IFNα-treated), and the nuclear transport inhibitor Orf6 (pSTAT, IFNα-treated) (**Figure 3a, Data Set S4**). Overall, our assays recorded several known viral hits, despite differences in cell lines, experimental conditions, and detection methods.

To further reveal trends of viral proteins that inhibit the innate immune pathways, we generated a separate list of 232 immunosuppressive proteins identified with a more permissive cutoff (**Figure 3a**, **Data Set S5**). We find large clusters of hits belonging to several flaviviral nonstructural proteins (25), such as 6 NS4B, 9 NS2a, and 5 NS4a sequences (**Figure 3c**). Other large clusters of protein hits include the aforementioned proteases and matrix proteins. We observe the same trend that viral proteins tend to be more pathway-specific inhibitors under our conditions (**Figure S4**). Notable proteins that broadly inhibited all three signaling pathways include SARS-Cov-2 Orf6, togavirin from Western equine encephalitis virus, and matrix protein from Jurona vesiculovirus. Additional coronaviral genes that inhibited immune pathways to a weaker extent include SARS-CoV Orf6 (26) and SARS-CoV-2 protease Nsp5.

### Mechanistic investigations into known, predicted, and new viral inhibitors

Protease activity is required for innate immune suppression by newly identified candidate proteases. 3C^pro^ from the picornaviral family are known to cleave various host factors in innate signaling pathways (27). For example, hepatovirus A (HAV) 3C^pro^ cleaves a bridging adaptor protein involved in IFN antiviral response in HEK293T cells (28). In cGAMP-stimulated BJ-5ta cells, we observed strong inhibition of IRF3-translocation when 3C^pro^ from parechovirus, hepatovirus A, and salivirus A were expressed (**Figure 4a**). 3C proteases from parechovirus and salivirus A have not been characterized to our knowledge. To demonstrate that the observed fold change is due to the proposed protease activity of viral proteins, we repeated our assay with the two candidate proteases and their corresponding catalytic cysteine variants. (**Figure 4a**). These mutants lost the inhibitory phenotype in our assay. Our results indicate that these other viral proteases, although weakly similar by sequence, are also capable of suppressing innate immune pathways.

**Figure 4:**
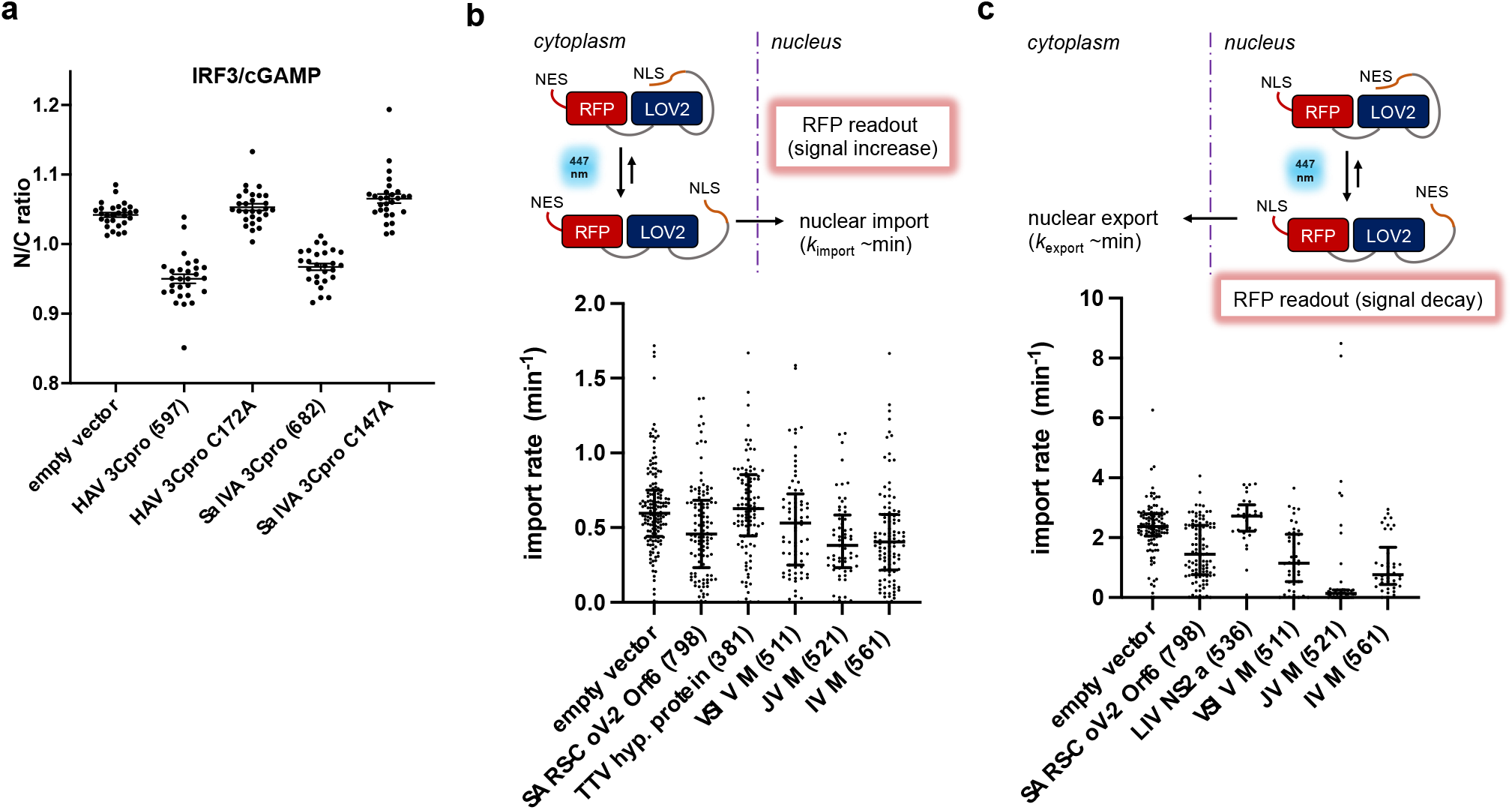
**(a)** Consolidated imaging results for nuclear translocation of IRF3 in cGAMP-stimulated BJ-5ta cells transfected with wild-type and variant viral genes. Cells transfected with hepatovirus A (HAV) 3C^pro^ and salivirus A (SalVA) 3C^pro^ exhibited lowered nuclear IRF3 intensity, while their corresponding active site mutants did not exhibit these effects. **(b), (c)** Matrix proteins from viruses in the *Rhabdoviridae* family inhibit nuclear import and export of RFP protein probe in U2OS cells. In the presence of 447nm light, a fusion protein with a LOV2 domain undergoes a conformational change that reveals either a nuclear localization signal (NLS, panel **b**) or nuclear export signal (NES, panel **c**) that increases or decreases nuclear RFP localization. Import and export rates were measured in single cells with a confocal microscope. These results demonstrated that M from vesicular stomatitis virus (VSIV), Isfahan virus (IV) and Jurona vesiculovirus (JV), which scored positive in our assay, inhibited nuclear import and export of proteins as expected. Torque teno virus 10 (TTV) hypothetical protein (381) and Louping ill virus (LIV) NS2a (536) were also tested in the assay as additional negative controls.

Newly identified matrix proteins (M) inhibit nuclear transport. Matrix proteins (M) encoded by the Rhabdovirus family were among our top hits. M from vesicular stomatitis virus (VSV) blocks host gene expression by binding to nuclear transport factors RAE 1 and Nup98 and inhibits poly(A) mRNA export (29). We find that M from Isfahan virus (IRF3 and NF-κB, poly(I:C)-treated), Jurona vesiculovirus (IRF3, cGAMP-treated), and vesicular stomatitis Indiana virus (NF-κB, poly(I:C)-treated) inhibited translocation in our imaged-based assays. Based on this result, we hypothesized that M from Isfahan and Jurona vesiculovirus also block nuclear import and export. We tested these matrix proteins in an optogenetics-based assay (30–32) that measures the rate of import and export of a fluorescent protein probe (**Figure 4b and c**). In this assay, U2OS cells stably expresses a photoactivatable nuclear transport signal fused to a red fluorescent protein with a constitutive nuclear export or import signal. For example, we measure a viral protein’s effect on nuclear import rate by expressing our viral proteins in U2OS cells that stably express a photoactivatable nuclear import sequence fused to a cytoplasmic RFP protein (**Figure 4b**). We find that the SARS-Cov-2 Orf6 (33), VSV M protein, and the uncharacterized rhabdoviral M proteins impaired bi-directional transport, consistent with blocking nuclear pores via interactions with Nup98.

We also identified several viral genes in which no immune inhibitory effects have been attributed to the best of our knowledge. We found 38 and 115 such genes under stringent and permissive cutoffs respectively (**Data Set 4 and 5**). Many of these proteins serve alternative functions in an infection, such as nucleoproteins, glycoproteins, capsid proteins, and matrix proteins. For example, we identified capsid proteins (Pfam accession: PF02956.15) from four different strains of torque teno virus that inhibited the STING and JAK/STAT pathways (**Data Set S5**). Two paramyxoviral glycoproteins (Pfam accession: PF00523) encoded by Hendra virus and human respirovirus 3 inhibited the TLR3 pathway.

Two proteins with no previously known function are a hypothetical protein with an intrinsically disordered domain encoded in torque teno virus 10 (TTV10) (NCBI accession: YP_003587850) and an intrinsically disordered protein from human respirovirus 3 (NCBI accession: NP_599250) (**Figure 5a**). TTV is reported to be a prevalent virus present in most humans, yet it is understudied along with other human anelloviruses due to difficult cultivating conditions (34). We tested these proteins in A549 Dual reporter cells (Invivogen ^®^) that report on the expression level of IRF translocation via a luciferase readout (**Figure 5b**). We observed that expression of TTV hypothetical protein and HRV disordered protein inhibited ISRE-driven gene expression when the cells were stimulated with poly(I:C). We next immunostained the streptavidin (strep)-tagged versions of the two proteins overexpressed in BJ-5ta cells and observed that TTV hypothetical protein is associated with the nuclear compartment (**Figure 5c**). However, this protein does not affect nuclear transport (**Figure 4b**, **Figure S5**) in our optogenetics assay. These data confirm the ability of our screen to identify traits of previously uncharacterized viral proteins.

**Figure 5:**
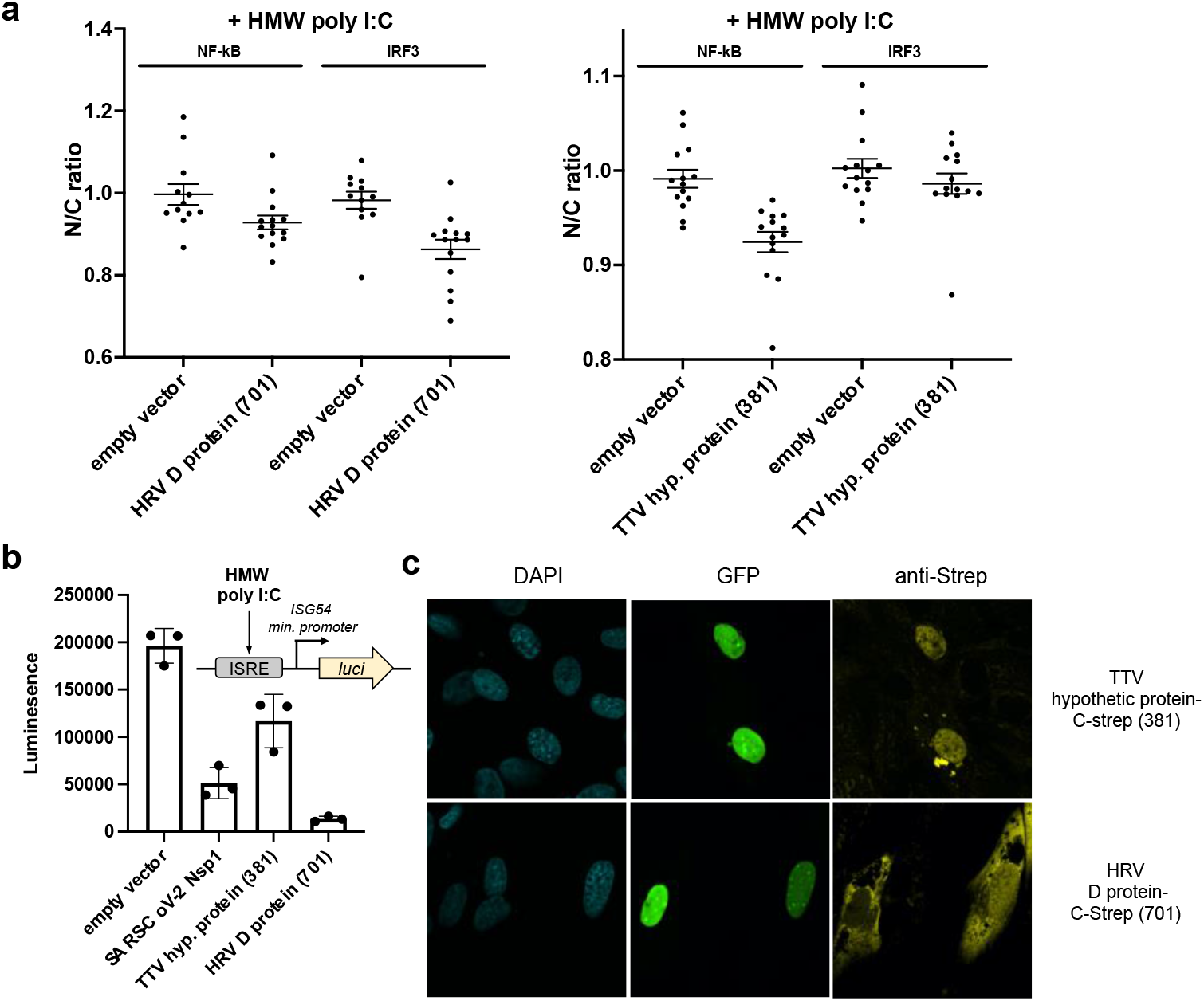
**(a)** High content imaging results for nuclear translocation of IRF3 when BJ-5ta cells express TTV hypothetical (hyp.) protein or human respirovirus (HRV) 3 D protein. Cells transfected with these genes exhibited lowered nuclear IRF3 intensity. **(b**) Reduced luminescence readout was observed for the TTV hypothetical protein and HRV D protein in A549 Dual (Invivogen^®^) cells when HMW poly(I:C) was used to stimulate the cells. ISRE = interferon stimulated response element. *luci* = gene encoding luciferase **C)** Cellular localization of the hypothetical TTV and HRV D proteins seen by immunofluorescence staining of C-terminal streptavidin-tagged proteins of interest expressed in BJ-5ta cells. SDS-PAGE of the overexpressed and purified viral proteins are shown in **Figure S5**.

## DISCUSSION

In this work, we characterized the ability of 800 viral proteins encoded by a diverse set of viruses to suppress host intracellular innate immune signaling pathways using a high-content, medium-throughput cell-based assay. We examined the nuclear localization of the transcription factors IRF3, NF-κB, and pSTAT1, which play key roles in the elaboration of the type I interferon response to viral infection, and which are major targets for inhibition by other viruses (35). To evaluate our assays, we undertook a bioinformatic approach that broadly searched databases of human, mammal, and insect infecting viruses in order to maximize the diversity of viruses we test. Our library was designed to contain known immune inhibitors as well as proteins that are related by sequence similarity or by containing homologous Pfam domains. We also included viral proteins with no predicted immunosuppressive function. Immune inhibitors within this class could be viral proteins that have an alternative function (*e.g*. capsid proteins) or in which no function has been assigned at all (*e.g*. hypothetical proteins). Altogether, this library allows us to benchmark the strengths and weaknesses of our assay, as well as provide an avenue for discovering new viral immune inhibitors. We note that this library will be available for other medium and high throughout assays, such as transcription-based reporter assays.

We developed an image-based screen to quantify the effects of viral proteins on nuclear translocation of various pro-inflammatory transcription factors. Viruses act on these pathways by a variety of mechanisms, such as specific inhibition of upstream signal transduction proteins, general inhibition of nuclear import, and enhancement of host factor degradation. Our assays involved transfection of a non-transformed cell line with expression vectors encoding each gene, followed by addition of a stimulator of innate immune signaling, and then immunofluorescence staining and automated image processing. This workflow allows for rapid testing of diverse viral genes and bypasses viral culturing. As such it allows us to test viral genes from any virus under low-containment conditions. Individual key signaling pathways were tested depending on the stimuli, and these assays allow us to probe effects of viral proteins on different stages of innate immune response. Some proteins may require concurrent viral infection for function and proper levels and thus appear as false negatives. However, our single protein assays concur with practices standard in the field (36, 37) and allow for a larger scale analysis as carried out here. We anticipate that these results would help steer further experimentation on the corresponding virus if possible.

Our assay reproduced numerous hits previously reported in the literature, such as poxin protein, rotavirus NS1, and numerous flaviviral protein clusters such as NS4B and NS2a, thereby confirming its value. While many hits were specific to one signaling axis, such as poxin and flaviviral NS2a, we did observe several hits that broadly blocked protein translocation, such as proteins that block nuclear transport. We focused on proteases from the ‘3C’ family and M proteins from rhabdoviruses as they were consistent hits, and we investigated their mechanisms *via* mutagenesis and nuclear import/export assays. These experiments confirmed that our predicted viral inhibitors identified from our assay exhibited the same immunosuppressive activities as previously characterized homologues. Furthermore, these results suggested our assay could facilitate mechanistic investigations and could also be optimized for assays to test for small molecule inhibitors of viral innate immune suppression.

Our screen also identified several hits that have not yet been assigned as immune inhibitors. Among these hits are capsid proteins from torque teno viruses and several paramyxoviruses. We also find paramyxoviral glycoproteins to be highly represented in our hit results. Viral proteins are known to be multi-functional, including structural proteins such as matrix or nucleocapsid proteins that may play a role in regulating innate immune responses (38, 39). We also identified hypothetical and uncharacterized proteins in which no functions have been elucidated to the best of our knowledge, such as the hypothetical protein encoded by torque teno virus 10. The results presented in our screen could guide future work to further dissect their mechanisms, potentially in the context of an infection. Our discovery of potential inhibitors from torque teno virus highlights the importance of culture-independent assays for functional characterization of proteins. Anelloviruses establish persistent infections in most healthy humans, and efforts to establish a reliable culture condition and to engineer these viruses for delivering payload are actively pursued (40). Assays similar to the ones presented in this study could forward our understanding and engineering of this viral family for translational applications.

Innate immune suppression may correlate with asymptomatic spread in addition to pathogenicity (41). By carrying out these experiments, we sought to provide tools for rapid, functional characterization of viral proteins in the era of metagenomics. This will further our capacity to link new viral sequences and function. Coupled with infection studies and/or independent secondary assays, our study could enable rapid testing of viral genes in assays for innate immune suppression, making it possible for early-stage evaluation of emerging and understudied viruses.

## MATERIALS AND METHODS

### Culturing BJ-5ta human fibroblast cell line

BJ-5ta cells were purchased from ATCC (CRL-4001) and cultured in the manufacturer recommended medium of 4:1 DMEM:M199 with 10% FBS (72% DMEM [ATCC 30-2002), 18% M199 [Thermo Fisher Scientific 11150059], 10% U.S Origin FBS (GenClone #25-514) with 10 μg/mL hygromycin B ([Invivogen ant-hg-1). Cells were cultured in T-25, T-75, and T-150 cell culture-treated flasks with vented caps (Corning) at 37 °C and 5% CO_2_. Cells were passaged every 3–4 days at 70-90% confluency to 30% confluency.

### Seeding BJ-5ta cells into 384-well plates

BJ-5ta cells were lifted from T-150 culture flasks using 0.25% trypsin (VWR 45000-664) for 5–10 min, then the trypsin was quenched with 2x volume of culture medium and transferred to 50 mL Falcon tubes. The cell suspension was centrifuged in a swinging-bucket rotor at 300g for 6 min at room temperature. The supernatant was discarded by aspiration and cells were resuspended in a small amount of culture volume and counted with a Bio-Rad TC20 Automated Cell Counter. Cells were diluted to 60,000/mL and 40 μL of diluted cell suspension was added from a reservoir to all wells except the outer row (*i.e*. rows B-O, columns 2-23 were used) of tissue-culture treated black CellCarrier-384 Ultra Microplates (Perkin Elmer 6057302) using a 12-channel electronic multichannel 200 μL pipettor [Sartorius]. Plates were then centrifuged at 200g for 4 min at room temperature and incubated overnight. Typically, about 2,000–5,000 cells per well are seeded, and about 200-800 are transfected as defined by expression of GFP.

### Co-transfection of virus gene plasmids and GFP into BJ-5ta cells

30–60 min prior to transfection BJ-5ta culture medium (4:1 DMEM:M199 with 10% FBS) was replaced with an equal volume of pre-warmed antibiotic-free transfection medium (4:1 DMEM:M199 with 20% FBS; 64% DMEM, 16% M199, 20% FBS). Transfection mixes were prepared according to manufacturer protocol (Lipofectamine 3000, ThermoFisher #L3000015) with final concentrations of 1 μg DNA, 4 μL GeneXPlus in 100 μL of Opti-MEM I Reduced-Serum Medium. Briefly, GeneXPlus [ATCC ACS-4004], plasmid DNA (200 ng/μL), and Opti-MEM I Reduced-Serum Medium (ThermoFisher #31985062) were warmed to room temperature and vortexed gently. Plasmid DNA was aliquoted into sterile microcentrifuge tubes at a ratio of 3:1 virus gene plasmid: GFP-containing plasmid. Opti-MEM was quickly mixed with GeneXPlus and the appropriate volume was added to each DNA aliquot and mixed briefly by gentle pipetting. GeneXPlus:DNA complexes were formed at room temperature for 15–20 min. Transfection mixtures were then added to each well at 10% final volume (4.4 μL transfection mixture was added to 40 μL transfection medium). Plates were centrifuged at 200 rcf, 4 min at room temperature to collect all transfection mixture into the medium and briefly mixed by tilting plate back and forth. Cells were incubated with transfection mixture at 37 °C, 5% CO_2_ for 24 h to allow DNA to enter cells. Transfection medium was then exchanged for fresh culture medium and cells were further incubated for another 24 h prior to stimulation with innate immune stimuli and fixation as described above.

### Homozygous knockout of cyclic GMP-AMP synthetase (cGAS) in BJ-5ta cells

CRISPR was used to introduce a frameshift mutation at position 13 of exon 1 of cyclic GMP-AMP synthetase (cGAS) in BJ-5ta cells. Three different Synthego-designed guide RNAs were each co-transfected with Cas9-containing plasmid (Synthego) into BJ-5ta cells (ATCC CRL-4001) using Lipofectamine 3000. After 48 h, samples were removed from each knockout pool for Inference of CRISPR Edits (ICE) analysis to assess gRNA efficiency, which was 1–6%. The knockout pool with 6% gRNA efficiency (gRNA 2) was diluted to a density of 0.5 cells/100uL and plated into 96-well plates for clonal expansion. Colonies grown from a single cell were visually identifiable after 3 weeks. After 8 weeks, the cGAS locus was sequenced in each clonal colony to identify colonies with homozygous indels. One homozygous knockout colony was identified from 20 screened colonies. Homozygous knockout in successful colonies was confirmed via Western Blot for cGAS protein.

### Stimulation of innate immune signaling

The (cGAS^−^)BJ-5ta cell line was used for these experiments. This cell line demonstrated a strongly reduced level of background innate immune signaling that otherwise resulted from introduction of transfecting DNA. In addition, cell transfection efficiency was improved relative to the parental BJ-5ta cells. Cells intended for stimulation with poly(I:C) HMW or liposome-encapsulated poly(I:C) LMW were primed 48 h in advance of stimulation by treating with interferon ɑ1 (Cell Signaling #8927) or interferon ɑ2b (PBL Assay Science #11100-1) at 50 ng/mL (final concentration in the well 5 ng/mL). 24 h after treatment with interferon, the cell medium was exchanged to remove external interferon from the cell environment. Different innate immune stimuli were applied to cell medium at 10% culture volume as follows: High molecular weight (HMW) poly(I:C) (Invivogen #tlrl-pic) at a concentration of 1mg/mL for 2 h (final concentration in the well 100 ug/mL) was used to stimulate TLR-3 activity by incubation at 37 °C, 5% CO_2_ for 2 h. 2’,3’-cyclic GMP-AMP (cGAMP) (Invivogen #tlrl-nacga23-5) at a concentration of 1 mg/mL (final concentration in the well 100 ug/mL) was used to stimulate STING pathway activity by incubation at 37 °C, 5% CO_2_ for 2 h. Interferon ɑ1 (Cell Signaling #8927) or ɑ2b (PBL Assay Science #11100-1) at a concentration of 50 ng/mL (final concentration in the well 5 ng/mL) was used to stimulate IFNAR activity by incubation at 37 °C, 5% CO_2_ for 45–50 min. Cell signaling was stopped by fixation as described below.

### Cell fixation and immunofluorescent staining

Cells were fixed with 15 μL of 16% methanol-free formaldehyde (ThermoFisher #28908) added directly to the 45 μL of cell medium in the wells for a final fixation solution of 4% formaldehyde. After 20–25 min incubation, the 4% formaldehyde solution was aspirated and the cells were washed three times with 60 μL PBS using an automated plate washer (BioTek EL406). Cells to be stained for phospho-STAT1 were further permeabilized with ice-cold 100% methanol (Sigma Aldritch #34860) and incubated at −20 °C for 10-15 min, then washed three times with 60 μL PBS using an automated plate washer. All primary and secondary antibodies were diluted 1:400 in PBS containing either 2.25% bovine serum albumin [Millipore Sigma #A2058] or 5% normal goat serum (Abcam #ab7481) for blocking and 0.15% Triton X-100 (Sigma Aldritch #T8787) for permeabilization. Fixed cells were stained with 40 μL diluted primary antibody solution overnight at 4 °C. Cells were then washed four times with 60 μL PBS using an automated plate washer and stained with 40 μL diluted secondary antibody solution with DAPI (ThermoFisher #D1306) added to a final concentration of 0.2 μg/mL. Cells were finally washed four times with 60 μL PBS using an automated plate washer and sealed using impermeable black plate seals. If not imaged immediately, fixed and stained cells were stored at 4 °C for a maximum of 4–7 days.

Cells treated with interferon ɑ were stained either for phospho-STAT1 (Cell Signaling Technology #9167) or for STAT1 (Cell Signaling Technology #14994). Cells treated with cGAMP or poly(I:C) HMW were stained simultaneously for IRF3 (Cell Signaling Technology #11904) and NFκB (Santa Cruz Biotechnology #sc-8008). IRF3, pSTAT1, and STAT1 primary antibodies were detected using an Alexa-Fluor 647-conjugated goat anti-rabbit IgG antibody (ThermoFisher #A21245). NF-κB primary antibody was detected using an Alexa-Fluor 568-conjugated donkey anti-mouse IgG secondary antibody (ThermoFisher #A10037).

### High-content imaging and image segmentation

Fluorescently stained plates were imaged on a PerkinElmer Operetta CLS High-Content Imaging System with a 20x, numerical aperture 0.75 objective. 20–25 sites were imaged in each well, covering 90–100% of the well. Each well was imaged for DAPI, GFP, and Alexa 647. Wells treated with cGAMP or poly(I:C) were also imaged for Alexa 568. Image segmentation was performed using Columbus software (PerkinElmer). Nuclear areas were identified with Columbus Method C based on the DAPI channel and cytoplasmic areas were assigned with Columbus Method D based on the Alexa 647 channel. Average intensities in the GFP, Alexa 647, and Alexa 568 (if applicable) channels were calculated for the cytosolic and nuclear areas of each computationally identified cell. Single-cell results were exported from Columbus in CSV format and can be requested from the authors.

### Data processing

Single-cell results were analyzed using a custom Python script which can be found at [Github URL TBD]. Briefly, nuclear objects identified by Columbus that correspond to cell debris and artifacts were eliminated based on nuclear morphology. For each transcription factor in a single cell, Nuclear Localization (Nucleus intensity / Cytosol intensity) and Total Cell intensity (Nucleus intensity + Cytosol intensity) were calculated. Within each well, cells were sorted into GFP positive (GFP+) or GFP negative (GFP-) (as a proxy for expression of virus protein) based on average Nucleus GFP intensity. The GFP positive or negative cutoff was set at twice the median Nuclear GFP intensity (the median being within the distribution of the more numerous GFP-negative cells). Within each well, the average Nuclear Localization or Total Cell intensity was calculated for the GFP+ or GFP-subsets of cells. Subsequently, for each well, the average Nuclear Localization or Total Cell intensity for the GFP+ cells was normalized to the corresponding average for GFP-cells to obtain a single normalized Mean Nuclear Localization or Mean Total Cell intensity.

Quality control was performed on a plate-by-plate basis as follows. If the mean of either of the two sets of controls containing no virus gene was outside the 20^th^–80^th^ percentiles of the plate data as a whole, the data for the aberrant control was discarded. If both no-gene controls were non-aberrant, the two sets of no-gene control data were combined for the following primary “fold change” calculation and normalization purposes. Additionally, for each individual set of 7 technical replicates, if any data point was more than 3 times the interquartile range higher than the 75^th^ percentile or lower than the 25^th^ percentile, it was removed from the analysis.

Within each plate, the mean and standard deviation of the 7 technical replicates of each virus gene/innate immune stimulus (one type of innate immune stimulus or mock stimulus per plate) combination were calculated. The fold change and corresponding statistical significance of each set of 7 wells for a given virus gene compared to the corresponding empty vector control were calculated. The raw values for the plots in **Figure 3** and **Figures S1–3** are attached in the Data Set S3 under the columns ‘logfold_T’ and ‘-log10pval’ (see Data Set descriptions).

### Blinded testing and analysis

For each protein, at least three different samples of the corresponding prepared DNA were given to a third party, who randomized and blinded them. The samples were then tested, analyzed, and the identity of each protein was assigned based on comparison to unblinded samples run concomitantly.

### Nuclear import and export assays

U2OS cell lines (engineered from HTB-96, ATCC) stably co-expressing Halo-H2A and NES-mCherry-LINuS or NLS-mCherry-LEXY were maintained in Dulbecco’s modified Eagle’s medium (DMEM, #10567022, Thermo Fisher) supplemented with 10% Fetal Bovine Serum (FBS, Thermo Fisher #A31605 or GenClone #25-514) at 37°C in a humidified atmosphere with 5% CO_2_. Cells were seeded at 10,000–15,000 cells per well in an 8-well chambered coverslip (#80826, ibidi) and grown in complete media for 1 day before co-transfection of the mammalian expression vectors encoding GFP and viral protein of interest using TransIT-2020 transfection reagent (#MIR5404, Mirus). After 24 h, the growth media was replaced with imaging media: low glucose (1g/L) DMEM without phenol red (#11054020, Thermo Fisher), supplemented with 10% FBS, 50 IU ml^-1^ penicillin and 50 μg ml^-1^ streptomycin, GlutaMAX^™^ Supplement (#35050061, Thermo Fisher). For nuclear staining, 500 nM JF646-HaloTag ligand (gift from Luke Lavis) was added in the imaging media. Image acquisition and kinetics measurements were performed as described previously (32).

### Transcriptional reporter assay from A549 Dual^®^ cells

A459 Dual^™^ (Invivogen) cells were maintained in F-12K medium (ATCC^®^ 30-2004^™^) supplemented with 10% heat inactivated Fetal Bovine Serum (FBS, Thermo Fisher Scientific #10082147), 10 μg/ml of blasticidin and 100 μg/ml of Zeocin^™^ at 37°C in a humidified atmosphere with 5% CO_2_. Cells were seeded into 96-well Flat Bottom TC treated culture plates (VWR^®^ 10062-900) at a density of 15,000 cells per well in F-12K medium with 10% FBS for 24 h. The next day, plasmids containing viral genes of interest, or empty vector control, were transfected using GeneXPlus according to manufacturer protocol. The transfected cells were allowed to grow and express the desired proteins for 48 h before high molecular weight poly(I:C) was added to the cells at a final concentration of 6 μg/mL. The cells were stimulated for 24 h at 37°C in a humidified atmosphere with 5% CO_2_. To measure IRF induction, 20 μL of the cell media supernatant was added to 50 μL of QUANTI-Luc^™^ assay solution (Invivogen #rep-qlc1) in a white 96-well microplate (Greiner, #655074), and luminescence was immediately measured using a BioTek UV-Vis spectrophotometer.

### Immunostaining and purification of overexpressed viral proteins

For immunostaining, BJ-5ta cells were seeded in 24-well glass-bottom culture plates (Cellvis P24-1.5H-N) at a density of 30,000 cells/well and allowed to grow at 37°C in a humidified atmosphere with 5% CO_2_ for 24 h. The next day, plasmids encoding torque teno virus hypothetical protein or human respirovirus 3 D protein were transfected into the cells using GeneXPlus according to manufacturer’s protocol. The transfected cells were allowed to grow for another 48 h. The cells were fixed with 16% methanol-free formaldehyde (ThermoFisher #28908) added directly to the cell medium in the wells for a final fixation solution of 4% formaldehyde. After 20–25 min incubation, the 4% formaldehyde solution was aspirated, and the cells were washed three times with 100 μL of PBS manually. Cells were stained for the streptavidin peptide using anti-Strep-tag II antibody (Abcam, #ab76949) diluted 1:2000 in PBS supplemented with 5% goat normal serum and 1% Triton-X. After incubating overnight at 4 °C, the cells were washed three times with 100 μL of PBS manually and stained with diluted secondary antibody solution (Alexa-Fluor 568-conjugated goat anti-rabbit IgG antibody) with DAPI (ThermoFisher #A-11011). After incubating for 1 h at room temperature, cells were finally washed four times with 100 μL PBS manually and imaged.

For testing protein expression, 1 x 10^6^ HEK293T cells were seeded in 150 mm cell culture dish and allowed to grow at 37 °C in a humidified atmosphere with 5% CO_2_ for 24 h. The next day, plasmids encoding empty vector, torque teno virus hypothetical protein or human respirovirus 3 D protein were transfected into the cells using TransIT-LT1 (Mirus Bio) according to manufacturer’s protocol. After 48 h, the cells were lifted with a cell lifter, washed with ice cold PBS, and the pellet was frozen at −80 °C at least 15 min before protein purification. To purify the proteins, the frozen cell pellet was allowed to thaw on ice for 30 min. 10 mL of lysis buffer (IBA LifeSciences Buffer W supplemented with 0.5% NP-40 substitute and one tablet of protease inhibitor (cOmplete^™^ EDTA-free Protease Inhibitor Cocktail, Sigma Millipore) was added to the thawed pellet. The mixture was passed through a syringe needle ~ 30 times before centrifugation at 12,000 rpm at 4 °C. The supernatant was passed through a column packed with Strep-Tactin^®^ resin (IBA LifeSciences #2-1208-010) and washed with IBA LifeSciences Buffer W (NC0612462, 3 x 10mL). The bound proteins were eluted from the column with 10 mL of IBA LifeSciences elution buffer and concentrated with a centrifugal concentrator. The concentrated soluble fraction was analyzed with SDS-PAGE or western blot using standard methods.

## Supporting information

Ng Data Set S1

Ng Data Set S2

Ng Data Set S3

Ng Data Set S4

Ng Data Set S5

## Data availability

All data and resources are available from the corresponding authors upon reasonable request.

## Acknowledgements

This work was supported by IARPA-FunGCAT Cooperative Agreement W911NF-17-2-0092. Tai L. Ng is an Open Philanthropy Awardee of the Life Sciences Research Foundation. Tae Yeon Yoo is supported by NIH/NIGMS postdoctoral fellowship F32GM131585. We thank Devin Burrill for editing the manuscript.

T.L.N., E.J.O., H.S.W., and T.M. performed plasmid purification, screens, data collection and analysis. T.L.N. and T.Y. performed optogenetic assay and T.Y. analyzed the results. Y. K. constructed the BJ-5ta cGAS knockout strain and expression vector design. E.J.O, P.D.K., P.M., T.J.M., T.Z.C., and J.Z. contributed to the screening methodology. N.J.R. and D.S.M. performed bioinformatics analysis. T.B. and C.M. contributed to the image processing. T.L.N., E.J.O., and N.J.R. wrote the manuscript. J.C.W. was primarily responsible for initial project conceptualization and subsequent direction with assistance from P.A.S., T.J.M., D.S.M, and J.I.G. P.A.S. was responsible for the overall project administration.

## Supplementary Materials

**Figure S1:** High content screen results for viral gene effects on IRF3 nuclear translocation in **(a)** high molecular weight (HMW) poly(I:C)-treated and **(b)** cGAMP-treated BJ-5ta cells. Volcano plots highlight hits for a total of 800 genes (including 195 coronavirus genes) with stringent cutoff lines in dashes. Cutoff values for log_2_fold change (log_2_FC) and p-values were set at 0.064 and 1.33 respectively in panel **a** and 0.067 and 1.33 respectively in panel **b**. These cutoffs were determined by comparing the data (left) to corresponding no treatment controls (right), which do not result in nuclear translocation of transcription factors.

**Figure S2:** High content screen results for viral gene effects on NF-κB nuclear translocation in **(a)** high molecular weight (HMW) poly(I:C)-treated and **(b)** cGAMP-treated BJ-5ta cells. Volcano plots highlight hits for a total of 800 genes (including 195 coronavirus genes) with stringent cutoff lines in dashes. Cutoff values for log_2_fold change (log_2_FC) and p-values were set at 0.06 and 1.33 respectively in panel **a** and 0.06 and 1.33 respectively in panel **b**. These cutoffs were determined by comparing the data (left) to corresponding no treatment controls (right), which do not result in nuclear translocation of transcription factors.

**Figure S3:** High content screen results for viral gene effects on pSTAT nuclear translocation in interferon (IFNα) treated BJ-5ta cells. Volcano plots highlight hits for a total of 800 genes (including 195 coronavirus genes) with stringent cutoff lines in dashes. Cutoff values for log_2_fold change (log_2_FC) and p-values were set at 0.085 and 1.33, respectively, as determined by comparing the data to the corresponding no treatment controls, which do not result in nuclear translocation of transcription factors. Parainfluenza 5 (PIV5) V protein is a positive control gene we use in most of our pSTAT translocation assays. We observe consistently that the protein inhibits pSTAT translocation in the presence of IFNα stimulus.

**Figure S4:** Venn diagram depicting distribution of 231 viral proteins across 3 assays that scored as positive hits under a more permissive cutoff (p<0.1). Most of the viral proteins tested were found to inhibit only one immune signaling axis of interest.

**Figure S5**: Commassie stained SDS-PAGE gels of **(a)** human respirovirus (HRV) 3 D protein (NCBI accession: NP_599250, expected size 45kDa) and **(b)** torque teno virus (TTV) hypothetical protein (NCBI accession: YP_003587850, expected band size 33kDa) overexpressed in HEK293T cells.

**DataSet S1:** List of 605 virus genes tested in this study. The spreadsheet also contains the annotations from our bioinformatics pipeline. Known immunosuppressors by GO ontology are listed as ‘1’ under column N (‘pos.control’). Predicted inhibitors based on sequence similarity are listed as ‘1’ under column O (‘pos.by.seq.identity’). Known immunosupressors by Pfam search are listed as ‘1’ under column P (‘pos.by.pfam’). Predicted immunosupressors by Pfam homology are listed as ‘1’ under column Q (‘pos.by.viral.architecture’). We also annotated viral proteins that contain protein domains with Pfam homology to human proteins, which are listed under columns R and S.

**DataSet S2:** List of 195 coronavirus genes tested in this study. The viruses tested are SARS-CoV-2, SARS-CoV, MERS-CoV, hCoV-229E, hCoV-NL63, hCoV-OC43, and hCoV-HKU1.

**DataSet S3:** Results for high content screen of 800 viral genes. Each tab corresponds to data from one screen testing one transcription factor and one stimulus. The data in **Figure 3** is derived from column B (‘logfold_T’) and column D (‘-log10pval). The data for no treatment controls in **Figures S1–3** is derived from column C (‘logfold_U’) and coliumn E (‘-log10pval_untreated’).

**DataSet S4:** List of hits using stringent cutoff at p < 0.05. The assay for which the viral protein scored positive is under Column B (‘Assay’).

**DataSet S5:** List of hits using permissive cutoff at p < 0.1. The assay for which the viral protein scored positive is under Column B (‘Assay’).

