## Supplementary figures and images for "High content screening and computational prediction reveal viral genes that suppress innate immune response"

### Figures S1-5

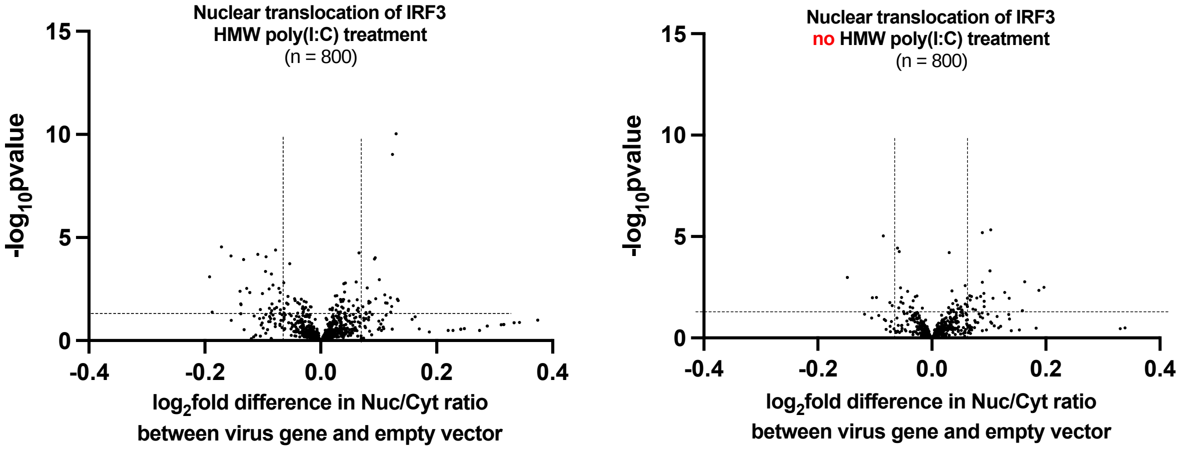

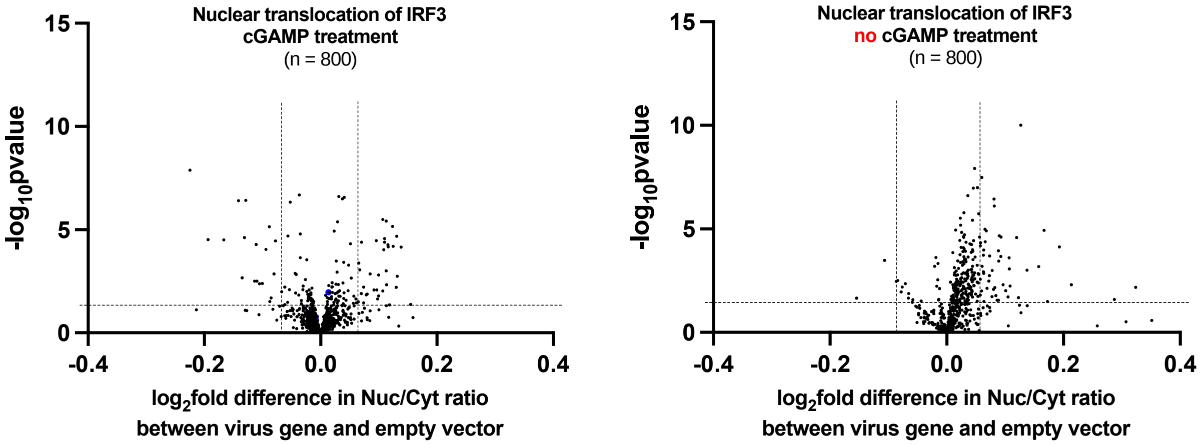

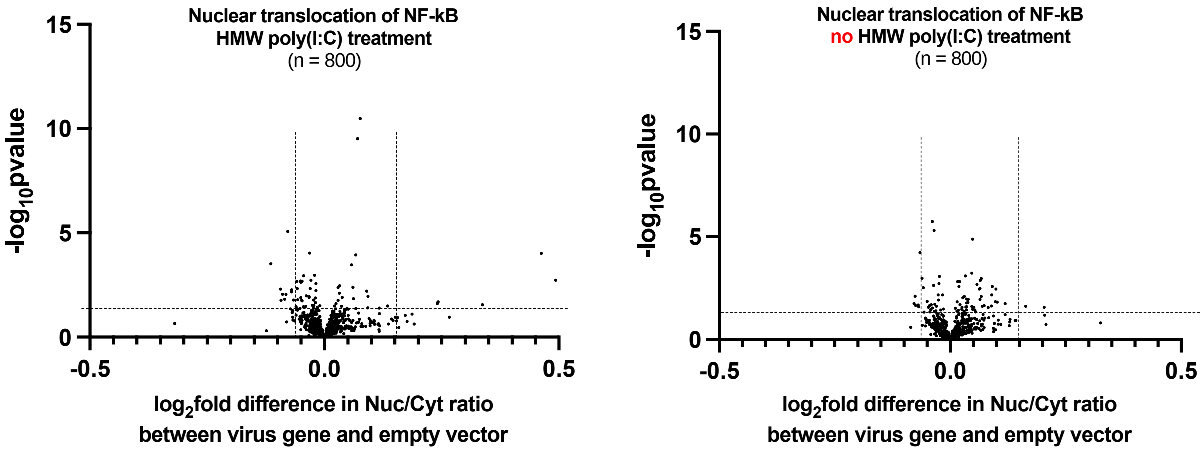

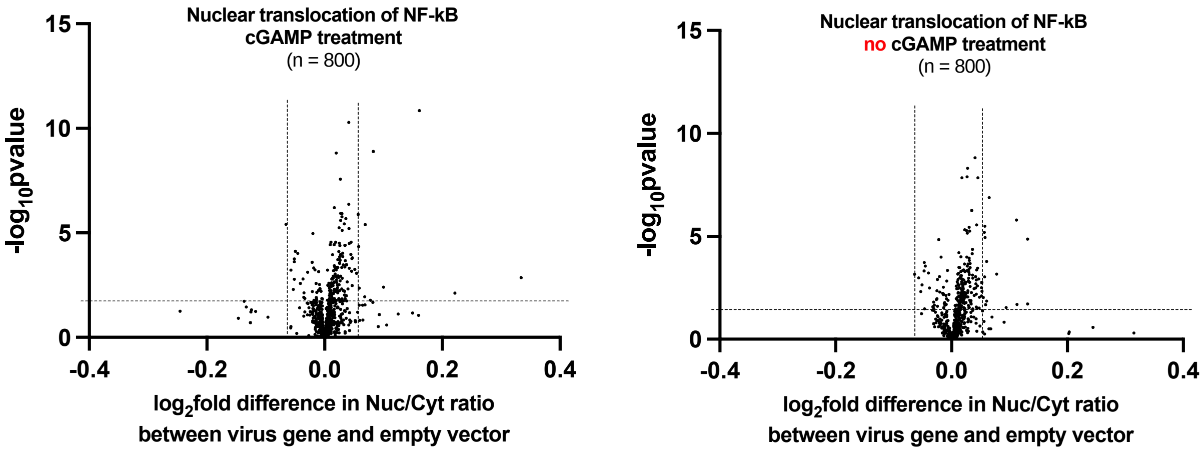

Figure S3

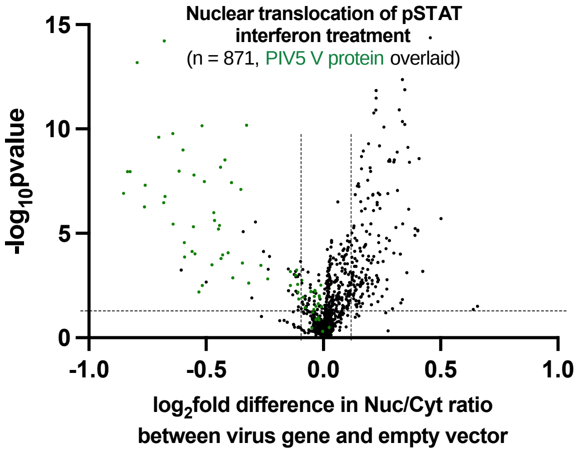


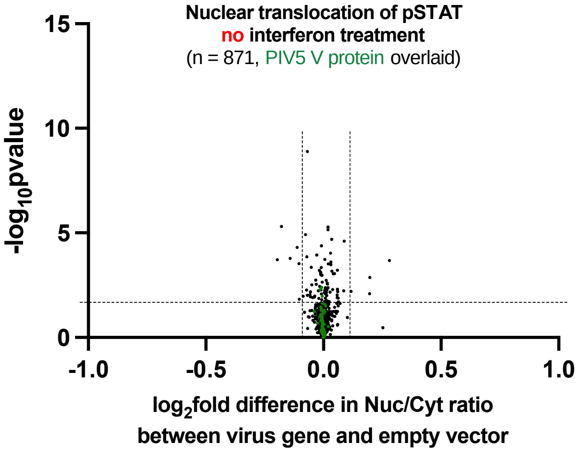


Figure S4

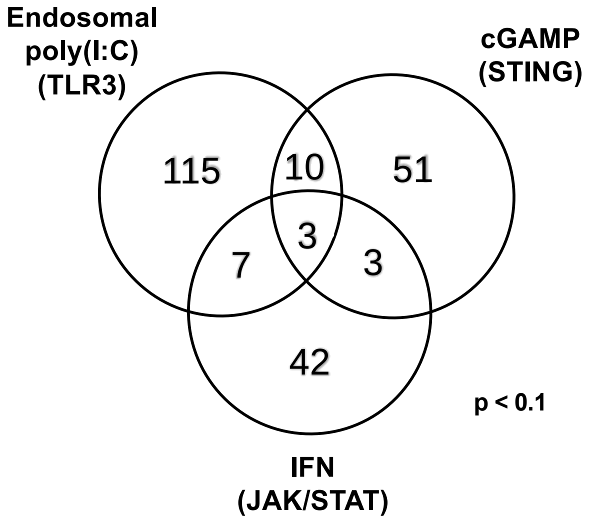

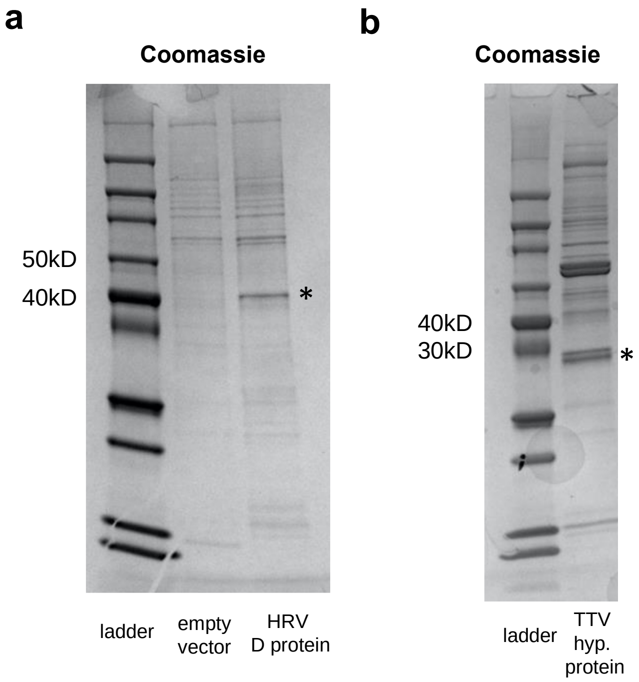
